# The optimal strategy of incompatible insect technique (IIT) using *Wolbachia* to control Malaria

**DOI:** 10.1101/2022.04.02.486813

**Authors:** Taiga Matsufuji, Sungrim Seirin-Lee

**Affiliations:** Graduate School of Integrated Sciences for Life, Hiroshima University, Higashi-hiroshima 739-8530, Japan; Institute for the Advanced Study of Human Biology(ASHBi), Kyoto University Institute for Advanced Study, Kyoto University, Kyoto 606-8315, Japan; JST CREST, 4-1-8 Honcho, Kawaguchi, Saitama 332-0012, Japan

**Keywords:** *Wolbachia*, Incompatible insect method, Malaria, Epidemic modeling

## Abstract

For decades, techniques to control vector population with low environmental impact have been widely explored in both field and theoretical studies. The incompatible insect method (IIT) using *Wolbachia*, based on cytoplasmic incompatibility, is a technique that *Wolbachia*-infected male mosquitoes are incapable of producing viable offspring after mating with wildtype female mosquitoes. While the IIT method experimentally ensured its effectiveness in several field works, the failure of female mosquito population control owing to the accidental contamination of *Wolbachia*-infected female mosquitoes has been a concern and an obstacle in implementing the IIT method in nature. In this study, we developed a population-based IIT mathematical model using cytoplasmic incompatibility and evaluated the effectiveness of the IIT method in scenarios where contamination was present or absent. In addition, by extending the model to assess the disease infection status of the human population with malaria, we evaluated the optimal release strategy and cost for successful disease control. Our study proves that IIT could be a promising method to control mosquito-borne diseases without perfect eradication of vector mosquito population regardless of contamination.

## 1 Introduction

Vector-borne infections account for more than 17% of all infectious diseases and cause more than 700,000 deaths annually. Malaria is a vector-borne disease transmitted by mosquitoes and is particularly devastating in the tropical and subtropical regions of the world (WHO 2019). The distribution of vector-borne infections is determined by a complex interaction of environmental and biological factors; therefore, the control of the vector population is critical for controlling the disease pandemic.

For decades, techniques for controlling the vector population with low environmental impact have been widely explored in both field and theoretical studies (Knipling 1955; Thomas et al. 2000; Phuc et al. 2007; Vreysen et al. 2007; Alphey et al. 2010; Lacroix and et al. 2012; Seirin-Lee et al. 2013a,b; Carvalho and et al. 2015; Natiello and Solari 2020). In particular, the strategy of releasing large numbers of male mosquitoes, which can reduce the mosquito population has been extensively applied. One classical control method is the sterile insect technique (SIT), in which males that have been sterilized using radiation do not produce eggs even after mating with wild females (Knipling 1955). The release of large numbers of sterile males results in a reduction of their reproductive output and potentially mosquito population abundance. As an improved method of SIT, transgenic technology, such as the release of insects carrying a dominant lethal (RIDL) (Thomas et al. 2000), has also been well explored for reducing mosquito populations (Lacroix and et al. 2012; Carvalho and et al. 2015). In RIDL, the released transgenic males are homozygous for a dominant lethal gene expressed in both male and female progeny that results from mating with wild-type insects. Genetically modified male mosquitoes are released and eggs laid by mating with wild females die in the larval stage, which consequently leads to the reduction of mosquito population. The control effectiveness for both classical SIT and RIDL has been verified by field experiments (Evans et al. 2019; Bouyer et al. 2020) and has been well studied in mathematical models so that possible control strategies have been examined and suggested under various situations (Phuc et al. 2007; Seirin-Lee et al. 2013a,b; Natiello and Solari 2020).

In recent years, the incompatible insect method (IIT) has attracted considerable attention as an alternative to the classical SIT method (Ritchie and et al. 2018). This approach relies on *Wolbachia*-infected male mosquitoes that are incapable of producing viable offspring after mating with wild-type females (Fig. 1A) (Laven 1967). *Wolbachia* is an intracellular symbiotic bacterium that infects a wide range of arthropods, including mosquitoes, and is thought to infect approximately 40% of the arthropod species in nature. Because it multiplies within the host cell, it cannot penetrate the sperm and is often localized in the ovaries of females; therefore, it is essentially transmitted from females to offspring via the egg (Werren et al. 2008). It causes four reproductive operations in host insects: feminization, in which a genetically male host is transformed into a female; parthenogenesis, in which a female gives birth to a female by herself; male killing, in which only male eggs are killed early in development; and cytoplasmic incompatibility, in which an infected male interferes with the reproduction of an uninfected female by causing abnormal cell division in the early embryonic stage of the egg resulting in failure to hatch. These four reproductive operations do not appear to be common and vary among hosts and *Wolbachia* lineages, whereas cytoplasmic incompatibility is most commonly found in many insect species (Werren et al. 2008).

**Figure 1:**
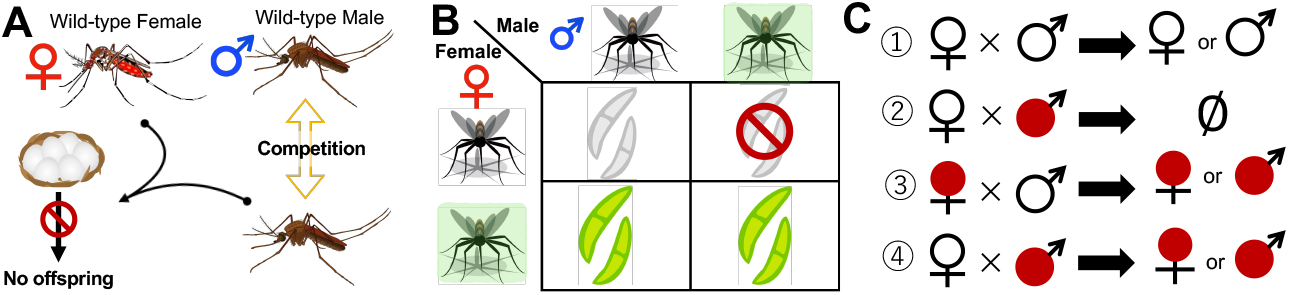
IIT images and combinations. (A) Schematic representation of IIT based on cytoplasmic incompatibility. The offspring do not hatch when a male mosquito infected with *Wolbachia* mates with a female mosquito of the wild type. (B) Results of mating between *Wolbachia*-infected (green shading) and uninfected (no shading) individuals. Infected females (bottom) always produced infected viable offspring, whereas uninfected females (top) produced uninfected viable offspring only when mating with uninfected males. In addition, mating between an uninfected female and an infected male result in non-viable offspring. (C) Four mating patterns are shown in (B). *Wolbachia*-infected/uninfected is marked by red/white color.

Recently, an IIT method based on cytoplasmic incompatibility was experimentally tested in Guangzhou, China (Zheng et al. 2019). In the experiment, male mosquitoes infected with *Wolbachia* were released and a high suppression rate of 83% − 94% in the population of wild mosquitoes was observed in the short term. On the other hand, if the released it Wolbachia-infected males are contaminated with *Wolbachia*-infected females, the IIT approach fails to sufficiently suppress the female mosquito population. This is because the eggs produced on mating of infected females hatch normally, regardless of the *Wolbachia* infectiveness of males (Fig. 1B and C). Namely, the *Wolbachia*-infected females leave offspring for the next generation and contamination of *Wolbachia*-infected females should be avoided for successful control in the IIT method.

Although the effectiveness of IIT using *Wolbachia* was prospectively shown in a field experiment, it is very difficult to confirm the effectiveness of a control strategy over a prolonged period in a wide region. Because a sustained release strategy of a large amount over a wide region requires high cost, the control strategy should be considered very carefully before implementation. Furthermore, if *Wolbachia*-infected females are contaminated by the release of *Wolbachia*-infected males, IIT is likely to fail. This implies that a strategy aimed at complete eradication of the mosquito population by IIT is not realistic in nature and a practically effective IIT strategy is necessary. To compensate for the limitations of field experiments, theoretical approaches using mathematical models have been applied to infer the long-term behavior of mosquito dynamics and test control strategies for control technologies (Phuc et al. 2007; White et al. 2010; Seirin-Lee et al. 2013a,b; Ewing et al. 2016; Kura et al. 2018; Natiello and Solari 2020), whereas the IIT method has been less explored theoretically.

In this study, we developed a mosquito population-based mathematical model describing the dynamics of *Wolbachia*-infected mosquitoes by using the IIT method based on cytoplasmic incompatibility. Using *in silico* experiments with the mathematical model, we evaluated the effectiveness of the IIT method in two scenarios of contamination of the *Wolbachia* female mosquitoes present or absent. In addition, we proposed an optimal release strategy for the elimination of female mosquito populations by introducing a release cost. Furthermore, we extended the model to assess the disease infection status of human populations rather than vector populations, especially for malaria. With the extended model, we primarily investigated the optimal release rate of the IIT strategy to reduce malaria infection rather than to target the extinction of female mosquito populations. Our study suggests that IIT could be a promising method to control mosquito-borne diseases without perfect eradication of mosquito populations.

## 2 Method

### 2.1 *Wolbachia* incompatible insect technique model (*Wolbachia* IIT model)

The life cycle of a mosquito is approximately 40 days. It takes approximately 15 days for the mosquito to reach adulthood via egg and larval stages (Carvajal-Lago et al. 2021). Thus, in this study, we assume that mosquito population growth proceeds via a stage-structured process and that density-dependent mortality acts on a pre-adult developmental stage. We choose the simple stage-structure model suggested by Dye (1984), in which the dynamics of wild-type mosquitoes are given to

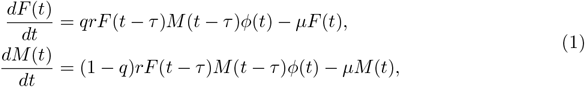

where *F* (*t*) and *M* (*t*) are the population densities of wild-type female and male mosquitoes, respectively, *r* is the survival rate from egg to adult, *q* is the sex ratio of female mosquitoes, *τ* is the growth time from egg to adult mosquitoes, and *µ* is the death rate. Function *ϕ* defines the density-dependent effect in the larval stage and is given as follows:

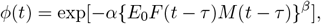

where *α* is a density-dependent coefficient, *E*_0_ is the egg production rate of adult mosquitoes without correction for density-independent survival between egg stage and adulthood and *β* is a parameter derived from fitting empirical data, as detailed in Dye (1984).

We extend the model (1) with *Wolbachia*-infected mosquitoes and develop an IIT model based on cytoplasmic incompatibility. In the cytoplasmic incompatibility technique, the offspring borne from mating between a wild female mosquito and a male mosquito infected by *Wolbachia* have chromosomal abnormalities and are not able to hatch normally (Fig. 1A). However, females infected by *Wolbachia* can lead to offspring regardless of the infectiveness of a male mosquito by *Wolbachia*, so that *Wolbachia* is inherited by the offspring (Fig. 1B and C). We denote the population densities of *Wolbachia*-infected female and male mosquitoes by *F*_*w*_(*t*) and *M*_*w*_(*t*), respectively. Subsequently, the IIT model, based on Fig. 1 is constructed as follows.

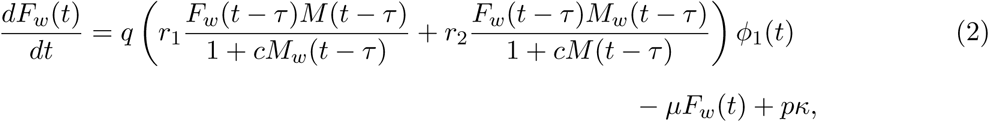

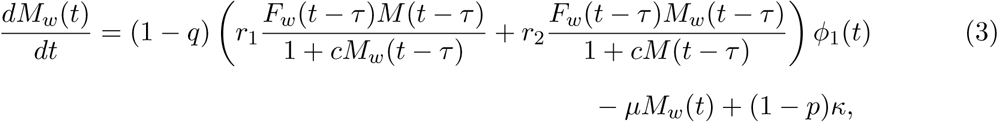

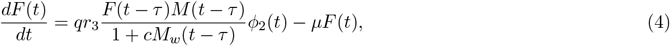

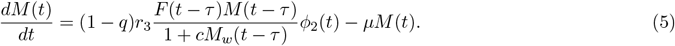

The first terms of Eqs. (2) and (3) are growth terms for mating mosquitoes and were modeled from ➂ and ➃ in Fig.1C, in which we assumed that wild-type male mosquitoes’ mate in proportion to their relative abundance (Knipling 1955; Phuc et al. 2007), and *c* (0 *< c* ≤ 1) represents the reduced mating competitive ability of *Wolbachia*-infected male or wild-type male mosquitoes. *r*_1_, *r*_2_, and *r*_3_ are the survival rates from egg to adult.

As wild-type offsprings are reproduced only by mating between wild-type mosquitoes (➀ in Fig.1C), the models for wild-type female and male mosquitoes are given in the basic model of Eq. (1) with adding the term mating competitive ability with *Wolbachia*-infected males. The density-dependent functions, *ϕ*_1_(*t*) and *ϕ*_2_(*t*), are given to

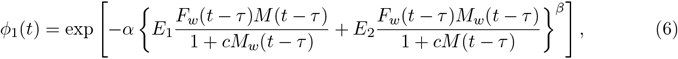

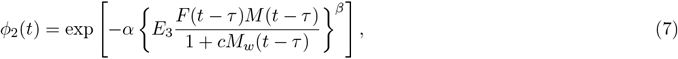

where *E*_1_, *E*_2_, and *E*_3_ are egg production rates for each mating combination. *α* and *β* are positive constants that determine the density-dependent effects.

The terminal terms in Eq. (2) and (3) are the release policies for the *Wolbachia*-infected mosquitoes and we assume a constant daily release of a fixed amount of mosquitoes and define it by a relative equilibrium density of wild-type female mosquitoes,

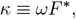

where *ω* is the release rate. *p* (0 ≤ *p <* 1) is the contamination ratio of the *Wolbachia*-infected female mosquitoes in the release policy. Note that the effect of reducing the number of offsprings using the IIT method (➁ in Fig.1C) is not explicitly included in our IIT model (2)-(5) because there is not hatching of eggs.

### 2.2 Application of the *Wolbachia* IIT model to Malaria (Malaria-IIT model)

In general, the control strategy of vector-borne diseases mainly focuses on the eradication of vector populations and aims at the extinction of vector insects. However, in the *Wolbachia* IIT model, such a goal may not be realistic because the *Wolbachia*-infected female produces offspring that can spread the disease. Thus, we examined the IIT strategy for malaria and considered the effectiveness of the IIT method in reducing infected cases. For this purpose, we combined a malaria transmission model comprising infected humans and mosquitoes.

A model to understand the dynamics of malaria transmission was first suggested by Ross (1911) and Lotka (1923). Subsequently, biological realism was given and plausible parameter values were estimated by Macdonald (1957). Ruan et al. (2008) extended the Ross-Macdonald model with combining the effect of the incubation period of malaria and updated the feasibility of model. Thus, we chose the Ross-Macdonald malaria model including the incubation periods of malaria.

By defining the malaria-infected human population as *H*(*t*) and the malaria-infected female mosquito population as *V* (*t*), the delayed Ross-Macdonald model with incubation periods is given by

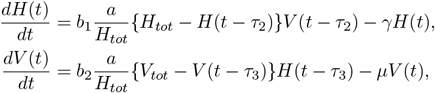

where *H*_*tot*_ is the total population density of humans, *V*_*tot*_ is the total population density of vector mosquitoes, *a/H*_*tot*_ is the average mosquito biting rate per individual, *b*_1_ is the transmission rate of malaria from infected female mosquitoes to uninfected human individuals through blood sucking, *b*_2_ is the transmission rate of malaria from infected individuals to uninfected female mosquitoes, *γ* is the recovery/mortality rate of infected patients, and *µ* is the mortality rate of malaria-infected female mosquitoes. *τ*_2_ and *τ*_3_ are the incubation periods for malaria in humans and mosquitoes, respectively.

Now, we incorporate the IIT strategy into the delayed Ross-Macdonald Malaria model by replacing the total population of vector mosquito *V*_*tot*_ with

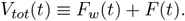

Note that only female mosquitoes bite humans; therefore, the male population is not included. Finally, we obtain the Malaria-IIT model as the following.

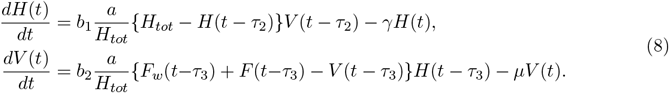

### 2.3 Parameters and initial conditions

We first assume that the sex ratio of mosquitoes is 1:1 (namely, *q* = 0.5) because cytoplasmic incompatibility does not critically change the sex ratio. For other parameter values, we chose those from previous studies that were estimated through experiments or mathematical verification. The detailed parameter set and references are shown in Table 1.

**Table 1:**
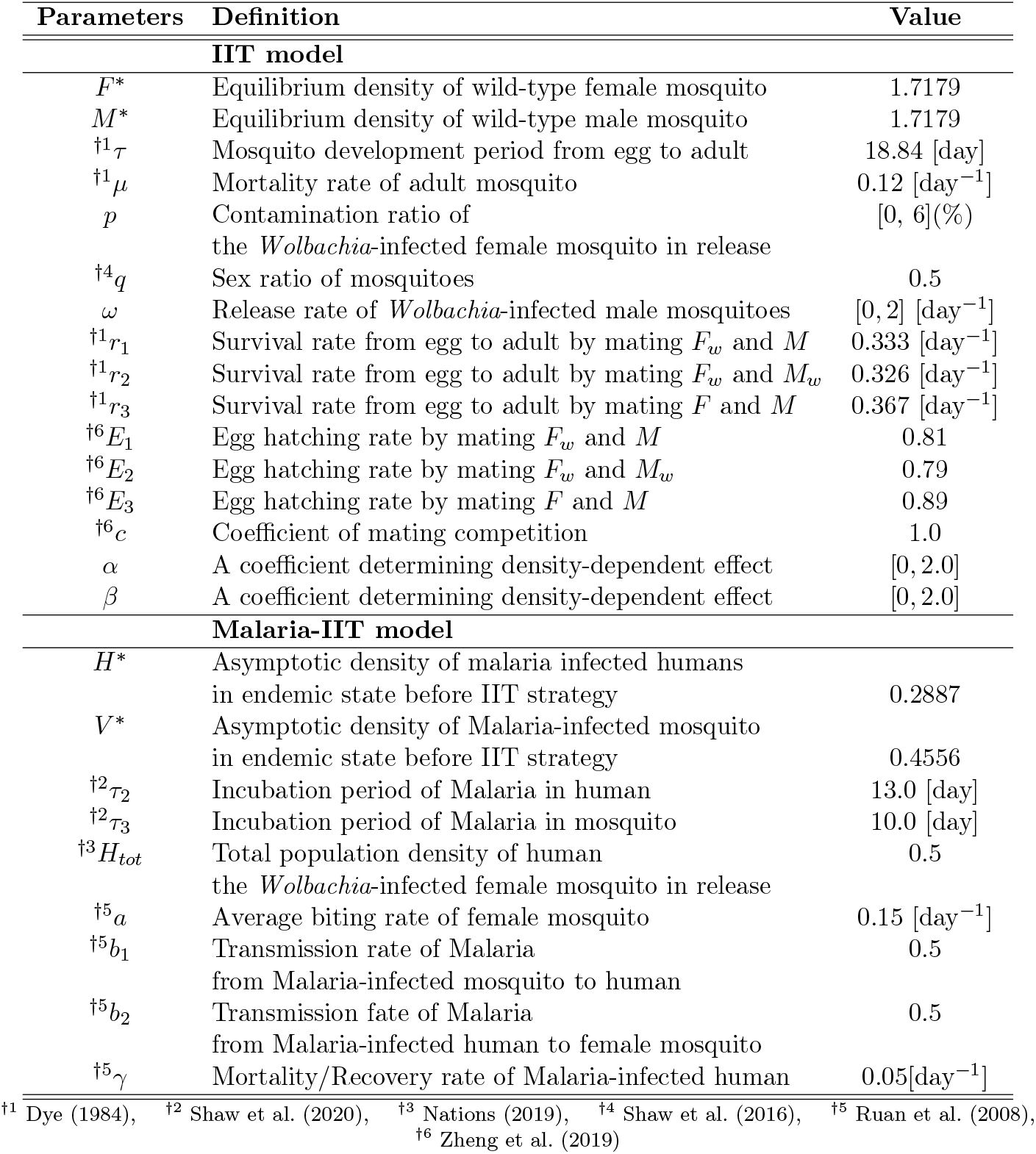
Representative Parameter set.

However, the parameters determining the effect of density dependence (*α* and *β*) in Eqs. (6) and (7) are unknown, although they are likely to influence the model dynamics (Phuc et al. 2007). Thus, to determine the feasible values of *α* and *β*, we directly tested the wild-type mosquito model (1) over a wide range of *α* and *β* values. We found that model dynamics can be classified according to the number of wild-type mosquito equilibria. We first note that there exists either the case of a unique equilibrium (0, 0) (red region of Fig. 2A), or the case of three equilibria, including (0, 0) (blue and green regions in Fig. 2A). Furthermore, the equilibrium (0, 0) is always locally stable, but the three equilibrium cases give rise to bi-stable states when we assume *τ* = 0 in Eq. (1) (see the Supplementary Information for the detailed analysis). In addition, we found that the maximal equilibrium density of female mosquitoes 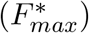 varies greatly depending on *α* and *β* in case of bistability. As the delayed differential equation likely causes oscillating dynamics, we presume that unrealistic oscillating dynamics may occur as 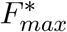 increases. Indeed, we often found oscillating dynamics in the bi-stable equilibria region of *α* and *β* (blue and green regions in Fig. 2A).

**Figure 2:**
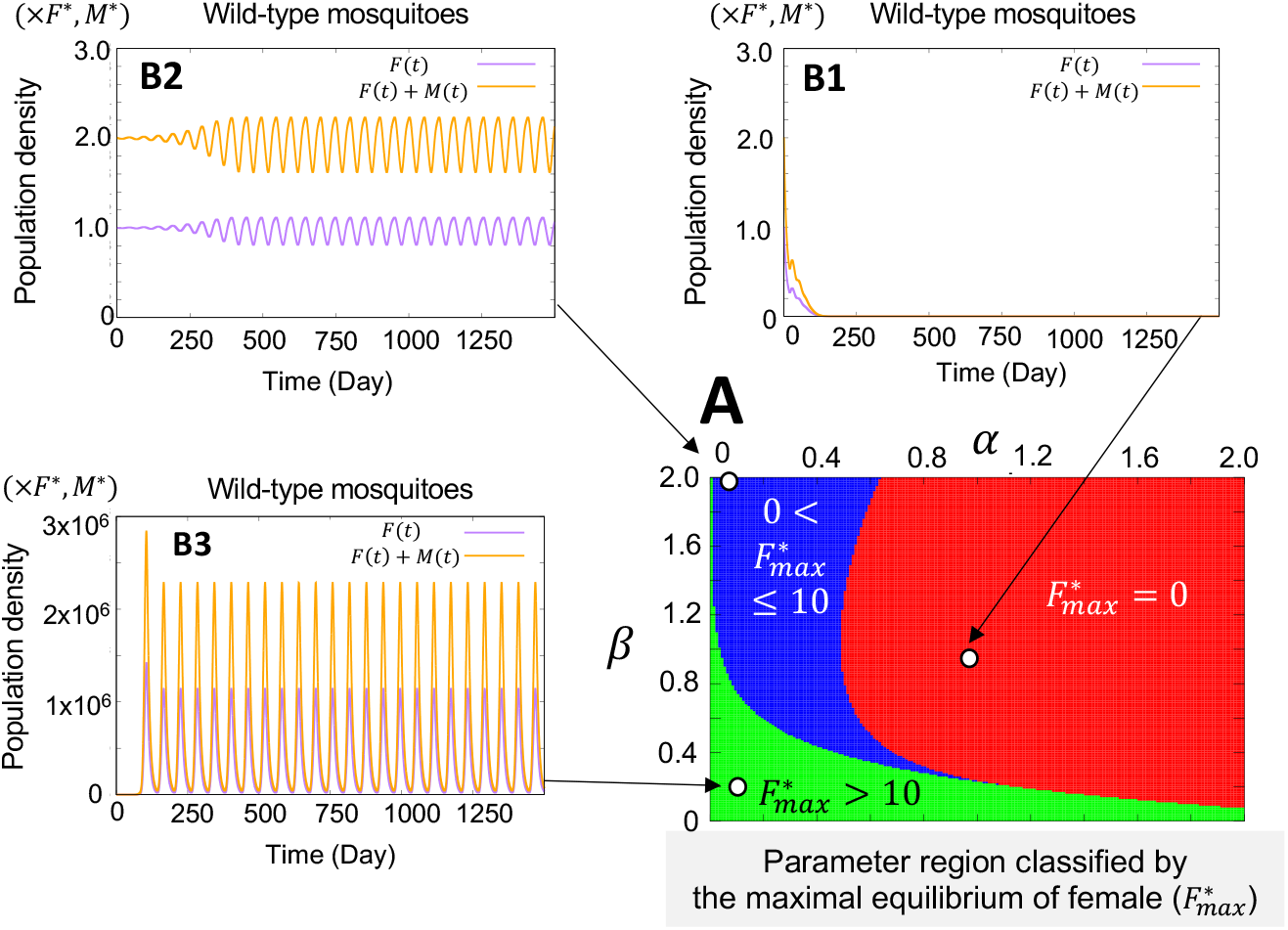
Parameter region and dynamics of the wild-type mosquitoes depending on (*α, β*). (A) The parameter region of (*α, β*) is classified according to the magnitude of the maximal equilibrium density 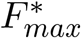. The red region corresponds to the case in which 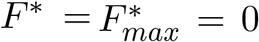 and (*F*\*, *M*\*) = (0, 0) becomes a unique equilibrium. The blue and green regions correspond to the three equilibrium cases and satisfy a bistable state. Namely, (*F*\*, *M*\*) = (0, 0), and 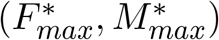 become stable equilibria. (B1-B3) The examples of the model (1) with *τ* = 18.84 days. (B1) shows the case of (*α, β*) = (1.0, 1.0), (B2) shows the case of (*α, β*) = (0.14, 2.0), and (B3) is the case of (*α, β*) = (0.1, 0.2).

Fig. 2 shows a representative result in which we investigated the parameter regions by 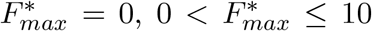, and 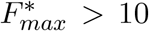. In the red region, the zero-equilibrium existed uniquely and mosquito populations were extinct when development period of mosquito(*τ* = 18.84 days) was chosen in the model (1) (Fig. 2B1). Thus, we excluded this parameter region. However, for the green and blue regions corresponding to the bistable region of equation (1) with *τ* = 0, the system showed oscillating dynamics when *τ* = 18.84 (Fig. 2B2, B3). Furthermore, the green region shows oscillating dynamics with a large amplitude of a dozen days period (Fig. 2B3), which is not a plausible dynamic for reflecting wild-type mosquitoes. In contrast, we found reliable parameter sets when 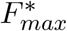 was not too large, where the mosquito populations oscillated with a low amplitude (Fig. 2B2). Thus, we chose *α* and *β* in a blue region of Fig. 2A.

Next, to define the initial conditions, we assumed that the *Wolbachia*-infected mosquitoes were non-existent in nature before release and the wild-type mosquitoes were at a stable non-zero equilibrium density. The details are given to

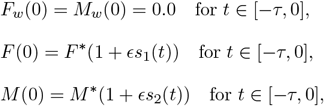

where *s*_1_(*t*) and *s*_2_(*t*) are chosen randomly in [0, 1], and *ϵ* = 0.01. Note that when the contamination ratio is zero (*p* = 0), the *Wolbachia*-infected female becomes *F*_*w*_(*t*) = 0 for all *t* ≥ 0 values.

Similarly, we assumed that the malaria-infected female mosquitoes and infected humans were in a state of co-existent equilibrium in the initial situation and then the IIT method was implemented. Thus, in the Malaria-IIT model, we adopted the following initial conditions:

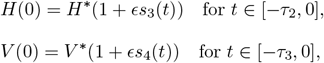

where *s*_3_(*t*) and *s*_4_(*t*) are chosen randomly in [0, 1], and *ϵ* = 0.01. *H*\* and *V* * are asymptotic densities obtained from equations (1) and (8) over a sufficiently long time.

## 3 Results

### 3.1 The effect of the *Wolbachia* IIT in the scenario without the contamination of the *Wolbachia*-infected female mosquitoes

Here, we investigated whether the *Wolbachia* IIT method is effective in eradicating the population of female mosquitoes assuming that contamination of the *Wolbachia*-infected female is absent in the release of the *Wolbachia*-infected male. Note that *F*_*w*_(*t*) ≡ 0, because *p* = 0 in this scenario (Fig. 3A; red lines).

**Figure 3:**
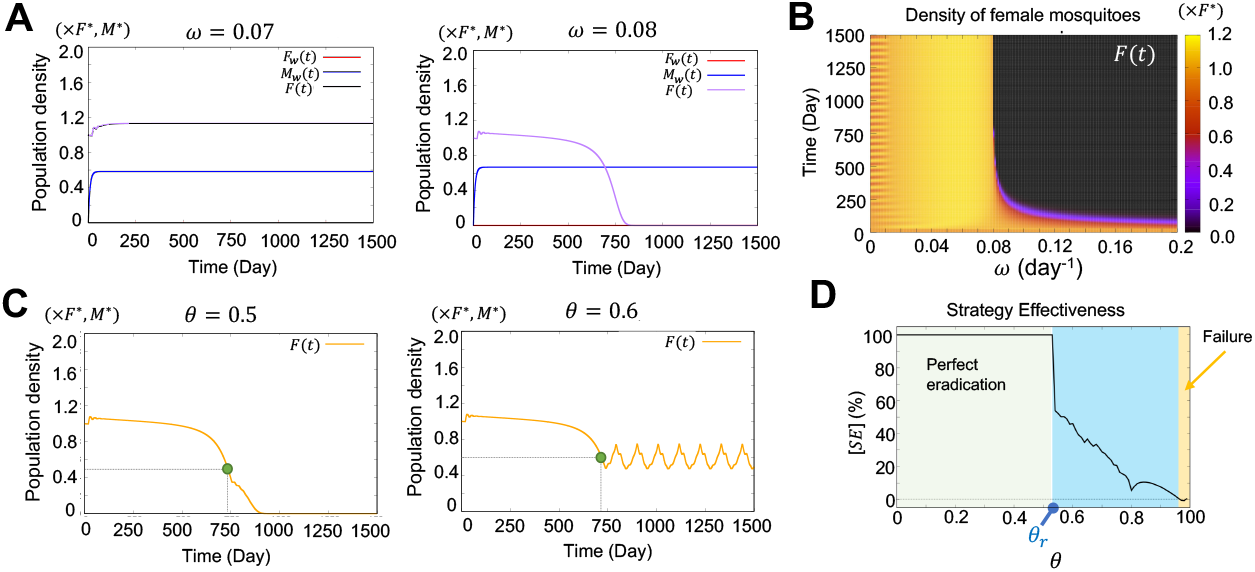
The effectiveness of the IIT method in the contamination absent scenario (*p* = 0). (A-B) The results of population dynamics by the release of the *Wolbachia*-infected male mosquito with varying release rates *ω*. As *p* = 0, *F*_*w*_(*t*) = 0 at all times. (C) Results obtained by choosing the release stop point (*θ*, Eq. (9)). The release was stopped at the 50% (left panel) and 60% point (right panel) of the initial density of the female mosquitoes. *ω* = 0.08. Green circles indicate the release stop points. (D) Strategy effectiveness calculated using Eqs. (10). The threshold value at which [*SE*] *<* 1 was approximately *θ*_*r*_ = 0.53. *ω* = 0.08.

We found that the IIT method was very effective when the release amount was sufficient (Fig. 3A and B). When the release rate (*ω*) was chosen to be less than 0.08, the wild-type female mosquitoes increased and the IIT method was ineffective (Fig. 3A, left panel). This was because the increase in male mosquitoes increased mating chances. Nonetheless, if the release rate was chosen to be a value larger than or equal to 0.08, the population of wild-type female mosquitoes dramatically decreased and the IIT method was very effective (Fig. 3A, right panel). This result indicates that our model is plausible for evaluating the effectiveness of the IIT method as shown in a field experiment (Mains et al. 2016; Zheng et al. 2019). In addition, when the release rate was greater than 0.08, the time for extinction of the female mosquito was not significantly dependent on the scale of the release rate (Fig. 3B). This suggests that if we choose a minimal release ratio to successfully eradicate female mosquitoes, the release cost can be significantly reduced.

Next, we examined whether continuous release is necessary to make female mosquitoes extinct. Thus, for a given release rate in the range of successful eradication, we stopped releasing the *Wolbachia*-infected male mosquitoes when the female mosquito population approached a certain level. We define release stop ratio (*θ*), satisfying

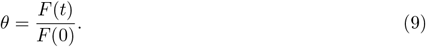

Specifically, *θ* refers to the population level of female mosquitoes compared to the initial population. We took *F* (0) = *F*\* as the initial equilibrium density of the wild-type female mosquito. Unexpectedly, we found that we could obtain successful eradication even when we stopped the release at *θ* = 0.5 and we need not continue the release of the *Wolbachia*-infected male mosquitoes (Fig. 3C). This result indicates that there exists an optimal stop point at which we can achieve successful eradication economically.

Thus, we investigated precisely how much the female mosquito population decreased depending on the release stop ratio *θ*. To explore it, we defined the strategy effectiveness by

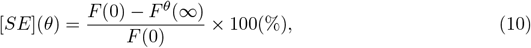

where *F* ^*θ*^(∞) is the asymptotic density of the female mosquito obtained over a sufficiently long time when the release of the *Wolbachia*-infected male mosquitoes is stopped at *θ*. In the simulation, *F* ^*θ*^(∞) was calculated as

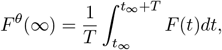

where *t*_*∞*_ is a sufficiently long time, and we set *T* = 280 (day) in the simulations. For strategy effectiveness, there exists a threshold value (*θ*_*r*_) of *θ*, and for *θ < θ*_*r*_, the perfect eradication of female mosquitoes is possible (Fig. 3C, green shadow region). Furthermore, we found that we can have a certain amount of effectiveness even when *θ > θ*_*r*_ (Fig. 3C, blue shadow region). However, when the release of the *Wolbachia*-infected male mosquitoes is stopped too early, the IIT method is ineffective and increases the density of female mosquitoes (Fig. 3C, yellow shaded region). Therefore, the IIT method can be a nature-friendly and cost-saving strategy to decrease vector mosquitoes effectively, at least, in the absence of contamination.

### 3.2 The effect of the *Wolbachia* IIT in a scenario with contamination of the *Wolbachia*-infected female mosquitoes

Here, we consider the *Wolbachia*-infected female mosquitoes are contaminated by the release of *Wolbachia*-infected male mosquitoes. We first tested the influence of contamination with the parameter set where the IIT method was successful in eradicating female mosquitoes when contamination of the *Wolbachia*-infected female was absent. As seen in Fig. 4A, the IIT method failed to eradicate female mosquitoes, even for a small contamination ratio (e.g., *p* = 0.1%) (Fig. 4B). This is because the IIT method is based on cytoplasmic incompatibility. The offspring could survive by mating the *Wolbachia*-infected females (Fig. 1C), so that the total population of female mosquitoes completely switched from wild-type to *Wolbachia*-infected female (Fig. 4A, second-line panels). However, the maximum density of female mosquitoes did not increase monotonically as the contamination ratio increased. For example, we have a lower density of female mosquitoes in the case of *p* = 2.0% than in those with *p* = 0.6%, 3.2% (Fig. 4A and B). Similarly, when we varied the release ratio (*ω*), we obtained a certain effectiveness for a range of release ratios (Fig. 4C and D). These results indicate that a perfect eradication is not possible when contamination occurs, while IIT is still effective in reducing the female mosquito density at a certain level by choosing an appropriate release rate with respect to a given contamination ratio.

**Figure 4:**
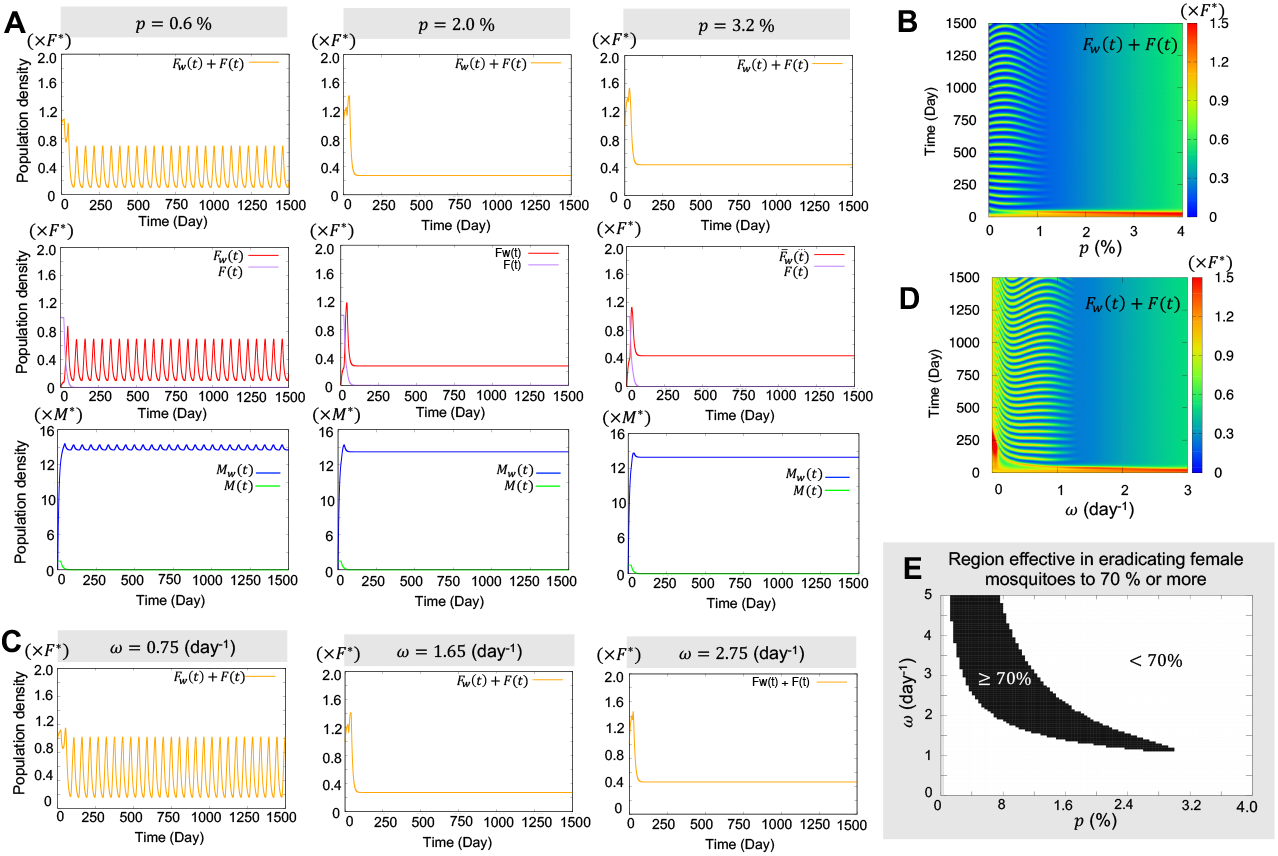
The effectiveness of the IIT method in the contamination present scenario (*p >* 0). (A) Population dynamics of mosquitoes with increasing contamination ratios (*p*). *ω* = 1.65. (B) Change in the maximum density of female mosquitoes as *p* increases. *ω* = 1.65 (C) Population dynamics of female mosquitoes with increasing release rate (*ω*). *p* = 2%. (D) Change in the maximal density of female mosquitoes with increasing release rate (*ω*). *p* = 2%. (E) The black region satisfies condition *F*_*w*_(*t*^*∞*^) + *F* (*t*^*∞*^) *<* 0.3*F*\*. *t*^*∞*^ was calculated for 7000 days after the total density of mosquitoes was less than 0.3*F*\*.

Thus, we next investigated a parameter region of the contamination ratio and release rate in which the total density of female mosquitoes can be eradicated to a certain level, for example, 70% or more (Fig. 4E). We found that there was a successful reduction in the release rate in some regions when the contamination ratio was less than approximately 3%, while the IIT became ineffective easily when the contamination ratio was high. Therefore, a high release rate with respect to a high contamination ratio is insufficient for reducing the density of female mosquitoes. This is because of the overabundance of *Wolbachia*-infected female mosquitoes caused by contamination in the large number of released mosquitoes. This proposes that the release rate should be carefully adjusted when the contamination exists.

### 3.3 Reduction effectiveness and release cost to reduce the female mosquitoes

In the previous section, we confirmed that contamination of *Wolbachia*-infected female mosquitoes is a fatal drawback of the IIT method based on cytoplasmic incompatibility. Nonetheless, the IIT method was still effective in reducing the density of female mosquitoes to a certain level. Naturally and technically, contamination cannot be controlled perfectly. Thus, we propose a policy to reduce the total population of female mosquitoes; however, it will not be a perfect eradication.

We investigated the effectiveness of IIT in reducing the density of female mosquitoes and the release cost required to reduce the female mosquito population to a certain level. To evaluate the reduction effectiveness of IIT, we calculated the reduction effectiveness [*RE*], defined by 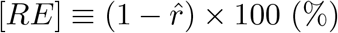 where

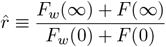

is the asymptotic density of the female mosquito by IIT control, and we define 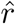 in 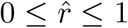. That is, the reduction effectiveness implies how much we can decrease the density of the female mosquito to a level of 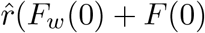 from the initial density, and is given by

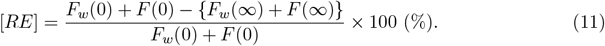

We investigated [*RE*] by varying the release rate (*ω*) and contamination ratio (*p*). In our model, we take *F*_*w*_(0) = 0, *F* (0) = *F*\*. In Fig. 5A, we first noticed that release effectiveness does not simply increase as the release rate increases. There exists a proper range of the release rate in which we can obtain a high release effectiveness with respect to a given contamination ratio (red-colored area in Fig. 5A), indicating that there exists an optimal release rate at which the effectiveness reaches its maximum.

**Figure 5:**
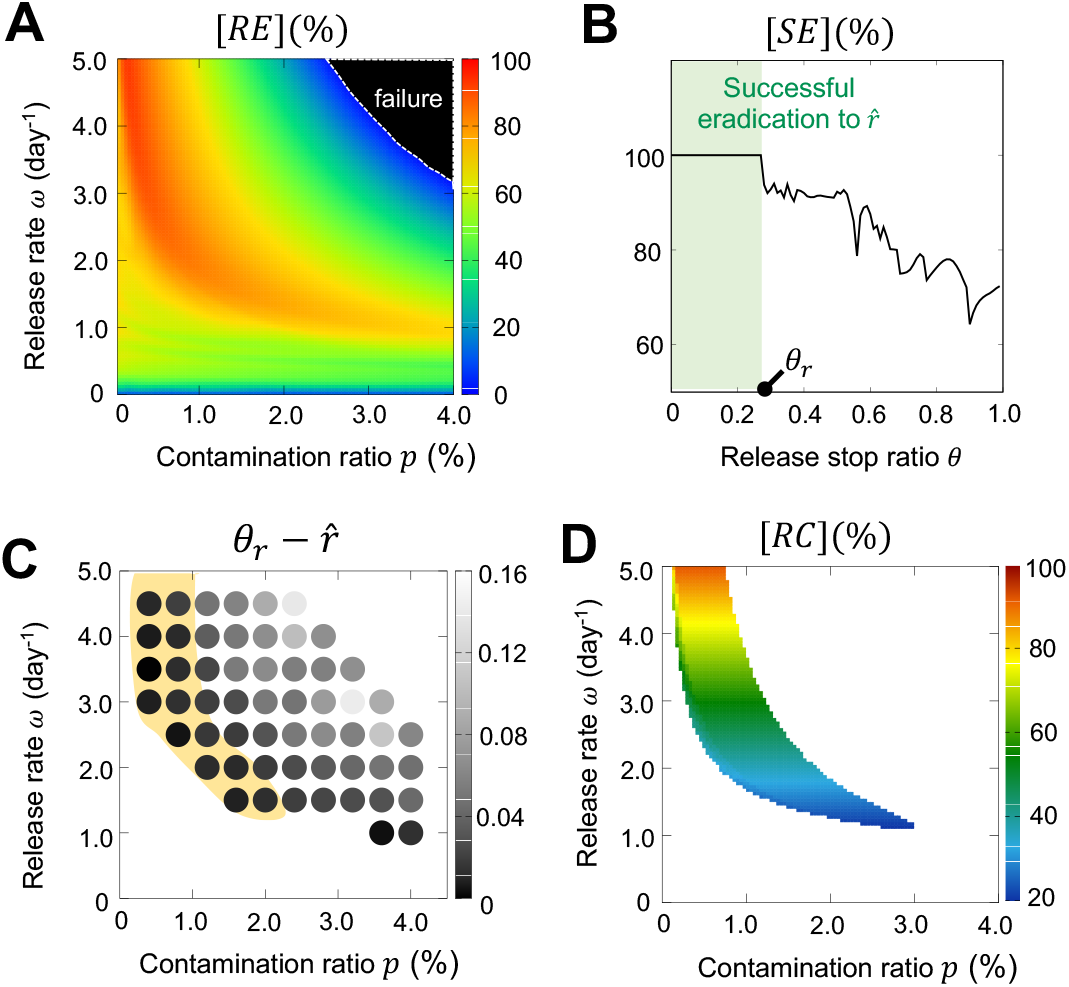
Release effectiveness and cost to reduce the female mosquitoes. (A) Release effectiveness calculated using Eq. (11), depending on the contamination ratio (*p*) and release rate (*ω*). The failure area is the parameter region in which the number of female mosquitoes increase more than the initial density. (B) Strategy effectiveness calculated by Eq. (13) for the case of *p* = 2%, *ω* = 1.67. The asymptotic density of the female mosquito was 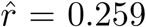, and the threshold value of the release stop ratio was *θ*_*r*_ = 0.27. (C) The effectiveness of the release stop strategy defined by the difference between the threshold value (*θ*_*r*_) of the release stop ratio and asymptotic density 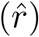 of the female mosquito. The yellow shadow region corresponds to the region of high release effectiveness (red region in (A)). The dark color implies that the *Wolbachia*-infected male mosquitoes must be released continuously until the density of female mosquitoes reaches the asymptotic density. (D) Release costs calculated using Eqs. (14) when [*RE*] ≥ 70% (i.e., 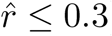).

Next, we further explore the strategy effectiveness function (13) to confirm whether there exists an effective release stop point:

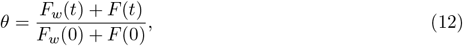

such that the total population of female mosquitoes reduces to a targeted density without extra release, namely, to an asymptotic density level 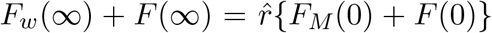, obtained after a sufficiently long time. The strategy effectiveness function is given as

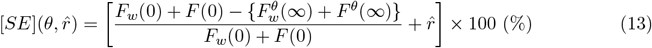

where 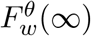 and *F* ^*θ*^(∞) are the asymptotic densities of the female mosquitoes after a sufficiently long time and when the release of the *Wolbachia*-infected male mosquitoes is stopped at *θ*. This measure implies that the reduction effectiveness reaches to 100 % when 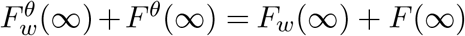.

We first found that a release stop ratio also exists in the scenario with contamination (Fig. 5B). Thus, we next explored the effectiveness of the release stop strategy in the case of imperfect eradication. We evaluated the measure of 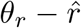, where *θ*_*r*_ is the threshold value of the release stop ratio. 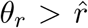 implies that we can have the same [*RE*] level without extra releases when the release is stopped at 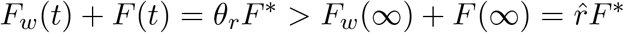. On the contrary, 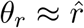 implies that the release stop strategy is not useful. We found that the release stop strategy is not useful especially in the parameter region where [*RE*] is high (the yellow shadow region in Fig. 5C). This result is in contrast with the scenario without contamination.

Because the release stop strategy was not effective, we finally evaluated the release cost defined by

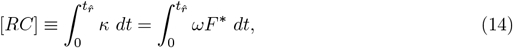

where 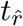 is a minimal time point satisfying 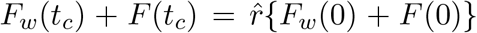. The release cost provides a measure of the number of *Wolbachia*-infected male mosquitoes that should be released to reduce the density of female mosquitoes to a certain level, namely, 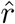. We calculated the release cost for the parameter region satisfying [*RE*] ≥ 70% and found that the release cost increases as the release ratio increases. This was because the time 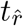 did not depend significantly on the release rates or the contamination ratios. Therefore, choosing a release rate where the [*RE*] becomes maximal in a given contamination is likely to be the optimal strategy to reduce the number of female mosquitoes without wasting costs.

### 3.4 Dynamics of Malaria infections after the *Wolbachia* IIT without contamination

The original goal of the transgenic insect strategy was to decrease the number of infected humans and control vector-borne diseases. Thus, we explored the effectiveness of the IIT method in decreasing the number of malaria cases. We first tested the dynamics of malaria patients and malaria-infected mosquitoes when *Wolbachia* mosquitoes were not introduced (Fig. 6A). The results showed that spread of malaria and its patients persisted as long as the population of female mosquitoes persisted. In contrast, the population of patients dramatically decreased once IIT was performed (Fig. 6B). With IIT, the number of patients decreased followed by a decrease in malaria-infected mosquitoes while there was a time gap in complete extinction. We also investigated how the release rate (*ω*) affects the effectiveness of the case and found that the IIT becomes effective when the release rate is larger than the same threshold release rate at which female mosquitoes become extinct (Fig. 6C, and Fig. 3B). This indicates that we can have successful extinction in both cases and in female mosquitoes.

**Figure 6:**
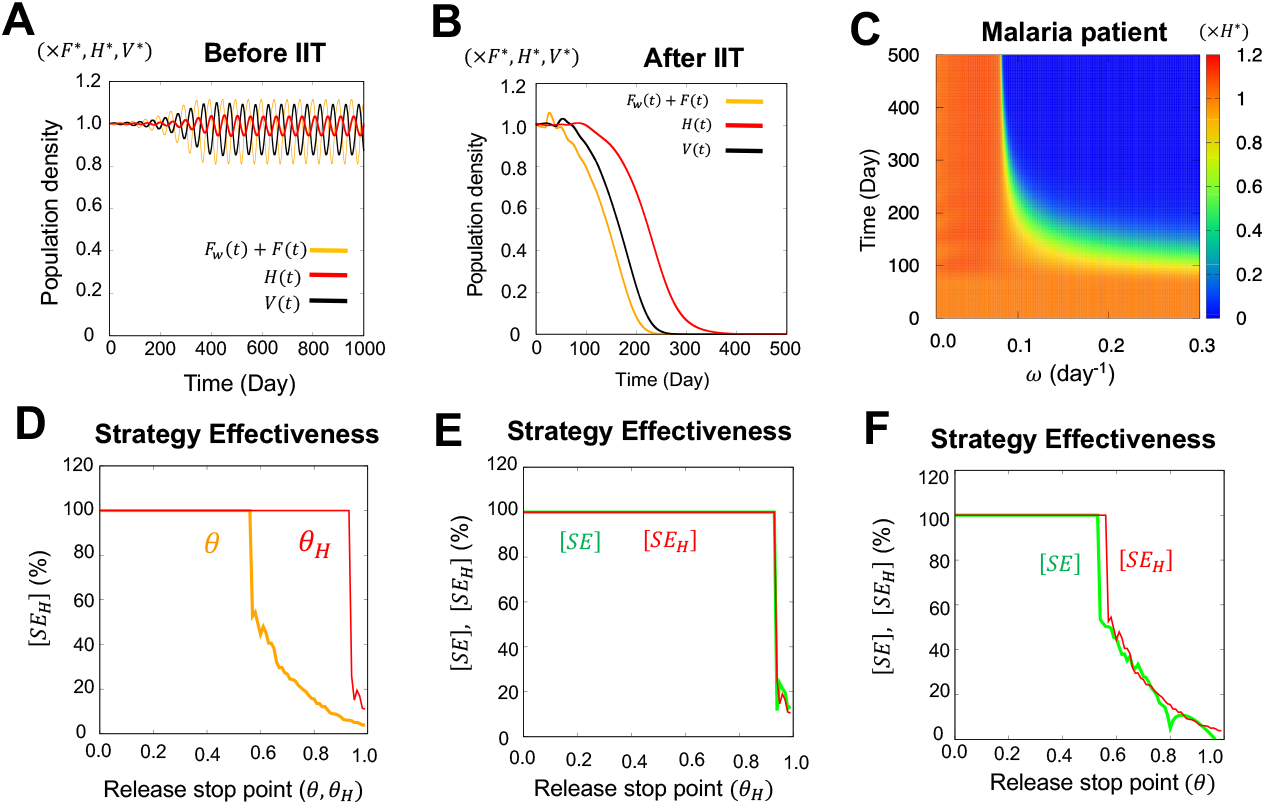
Malaria-IIT model and strategy effectiveness in the contamination absent scenario (*p* = 0). (A) Population dynamics of female mosquitoes (*F*_*w*_(*t*) + *F* (*t*)), Malaria-infected humans (*H*(*t*)), and Malaria-infected mosquitoes (*V* (*t*)) before conducting the IIT method. Model equations (1) and (8) were simulated. (B) Population dynamics after IIT. The model equations (2)-(5) and (8) were simulated with *ω* = 0.1. (C) Effect of the IIT method on release ratio *ω*. (D) Strategy effectiveness ([*SE*_*H*_]) with respect to release stop ratio *θ* (Eq. (12)) and *θ*_*H*_ (Eq. (15)). *ω* = 0.1 (E-F) Strategy effectiveness ([*SE*] and [*SE*_*H*_]) with respect to release stop ratios *θ*_*H*_ and *θ. ω* = 0.1

Next, we explored strategy effectiveness in a manner similar to that in Fig. 3D with respect to the release stop point (*θ*) for the female mosquitoes. In addition, we explored the release stop point (*θ*_*H*_) for an infected human such that

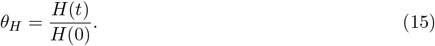

We also defined the strategy effectiveness based on the population of infected humans as follows:

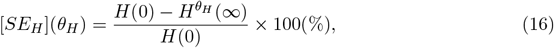

where 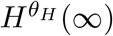 is the asymptotic density of the patients after a sufficiently long time and when the release of the *Wolbachia*-infected male mosquitoes is stopped at *θ*_*H*_. In the simulation, 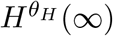 was calculated as

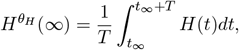

where *t*_*∞*_ is a sufficiently long time and *T* is taken as 280 days.

We first investigated the strategy effectiveness [*SE*_*H*_] with respect to the two release stop points, *θ* and *θ*_*H*_ in Fig. 6D, respectively. We found that both *θ* and *θ*_*H*_ can provide perfect effectiveness when *θ* ≤ 0.56 or *θ*_*H*_ ≤ 0.93. This suggests that we can control the pandemic without releasing the *Wolbachia*-infected mosquitoes until the female mosquitoes are completely eradicated or the large number of infected humans decrease.

Finally, to compare which strategy effectiveness should be chosen as a measure that incurs less release costs, we investigate [*SE*] and [*SE*_*H*_] with respect to both *θ*_*H*_ and *θ* (Fig. 6E and F). We found no notable difference between the effectiveness of the two strategies for both *θ*_*H*_ and *θ*, indicating that we may choose either release stop measures or strategy effectiveness measures at least under the scenario of no contamination.

### 3.5 The effectiveness of the *Wolbachia* IIT on malaria and optimal release cost under the scenario of contamination present

In the previous section, when *Wolbachia*-infected female mosquitoes were contaminated by the IIT method, perfect eradication was impossible; however, we obtained a certain level of strategy effectiveness depending on the release rate (*ω*) and contamination ratio (*p*). Thus, we examined the effectiveness of IIT in reducing the population of malaria-infected cases in the presence of contamination. We first investigated the dynamics of malaria cases by varying *ω* and *p* (Fig. 7A and B). Unexpectedly, we found that we can have a perfect extinction of malaria-infected cases, even though female mosquitoes were not extinct (Fig. 7A and B, the second panels, and Fig. 7E). However, once the female mosquitoes persist for more than a certain population level, the IIT method could not lead to a perfect eradication of malaria, although we still could have a certain level of effectiveness in reducing the population of malaria-infected cases (Fig. 7A and B, first and third panels, and Fig. 7C and D).

**Figure 7:**
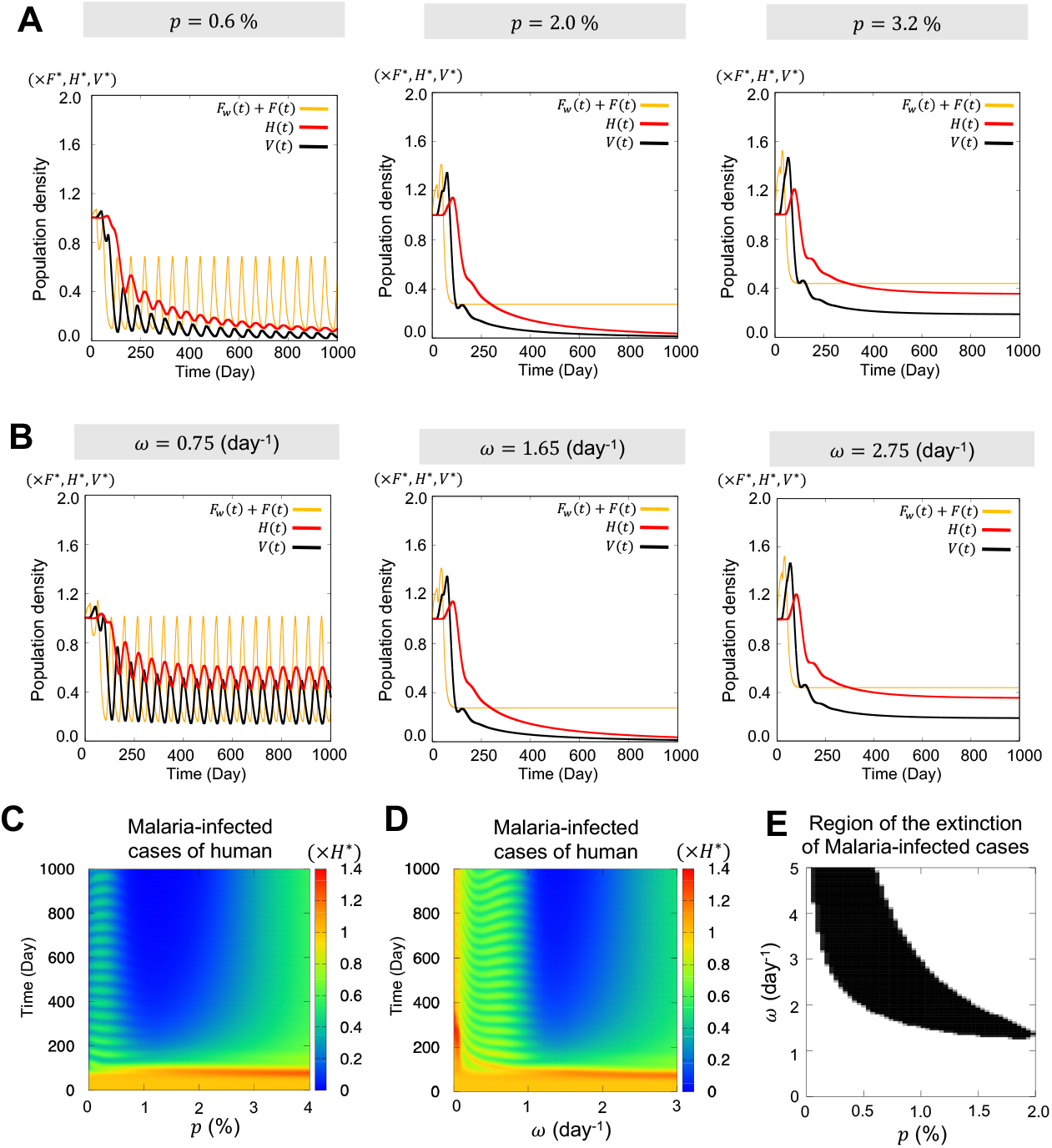
The effectiveness of the *Wolbachia* IIT under the scenario of contamination present. (A) Population dynamics of Malaria-infected female mosquitoes with varying contamination ratios (*p*). *ω* = 1.65 was selected. The parameters of the IIT model are the same as those in Fig. 4. (B) Population dynamics of malaria infected cases and female mosquitoes with varying release rates (*ω*). *p* = 2.0% was chosen. The parameters of the IIT model are the same as those in Fig. 4. (C-D) Population dynamics of Malaria-infected humans with varying contamination ratios and release rates. *ω* = 1.65 was fixed in (C) and *p* = 2.0% was fixed in (D). (E) Parameter region where malaria-infected cases are extinct. The region was calculated using the condition satisfying *H*(*t*)*/H*\* ≤ 0.001 for a sufficiently long time.

Finally, we calculated the release cost and explored the optimal release ratio values for a given contamination ratio. Because we were able to obtain a perfect eradication using the IIT method as shown in Fig. 7E, we first investigated the release cost, [*RC*_*H*_], defined by

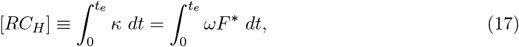

where *t*_*e*_ is the extinction time, that is, the minimal time point at which the malaria-infected cases of humans are extinct. To avoid additional calculations owing to numerical errors, we set *t*_*e*_ = min{*t*|*H*(*t*_*e*_) ≤ *εH*\* (*ε* ≪ 1)}.

The result of Fig. 8A shows that the release cost is not simply determined by the release rate monotonously and there exists an optimal release rate at which the release cost becomes minimal for a given contamination ratio. Thus, we further investigated where the minimal release cost was obtained in the range of the release rate shown in Fig. 8B. We found that the minimal release cost increased as the contamination ratio increased, whereas the optimal release rate satisfying the minimal release cost decreased. This indicates that when the contamination ratio is high, we must choose a lower release rate. On the other hand, a long time was required for cases to become extinct when we chose a low release rate, as shown in Fig. 8C. If the disease persists for a long time, the number of deaths will increase. Thus, when contamination is large, the policy of minimal release cost is not always the best.

**Figure 8:**
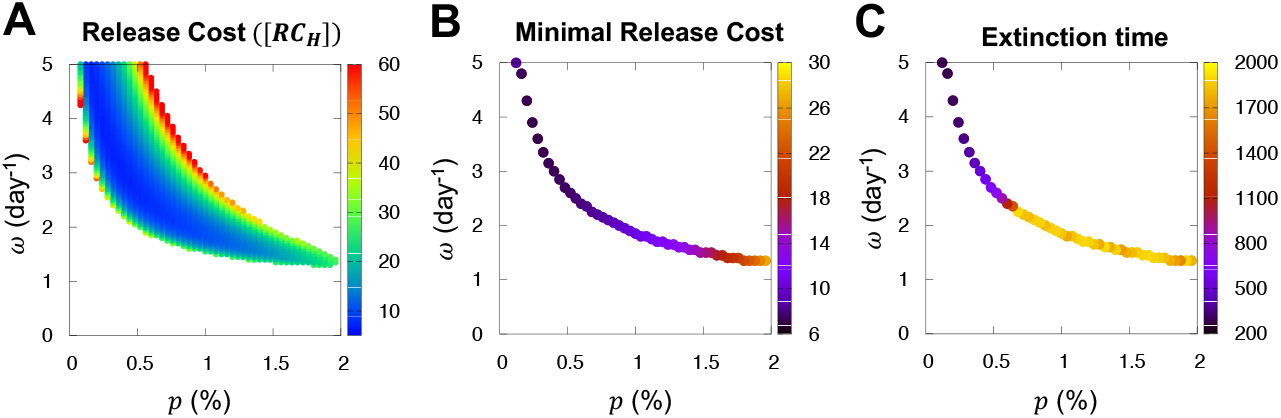
Release cost and optimal release ratio. (A) The release cost ([*RC*_*H*_]) given by Eq. (17) was calculated by varying the release rate(*ω*) and contamination ratio (*p*). (B) The minimal release cost plotted from (A) for a given contamination ratio. (C) Extinction time *t*_*e*_ plotted from (A) for a given contamination ratio.

Taken together, we concluded that the IIT method can be a very effective strategy to eradicate malaria but the optimal policy to reduce the cost should be carefully considered depending on the contamination ratio and extinction period of the cases.

## 4 Discussion

Incompatible insect technique (IIT) using maternally inherited endosymbiotic bacteria *Wolbachia* is considered a promising method that can avoid an influence on male mating competitiveness, survivability, and nature. Thus, the IIT method was chosen as an alternative to the sterile insect technique (SIT); in particular, it has been considered an effective method for the suppression of mosquito populations (Curtis et al. 1982). However, the contamination problem of the *Wolbachia*-infected female mosquito in the IIT method has been a concern because the *Wolbachia*-infected female mosquito leads to the replacement of the wild-type field mosquito population with the *Wolbachia*-infected mosquitoes, which results in the failure of disease control.

In this study, we developed a mathematical model based on the population dynamics of *Wolbachia*-infected mosquitoes and explored the extent to which the IIT method and the release strategy of *Wolbachia*-infected male mosquitoes can effectively suppress female mosquitoes in the two scenarios of the contamination being absent or present. As shown in previous field studies (Curtis et al. 1982; Mains et al. 2016; Zheng et al. 2019), the IIT method was highly effective in eradicating wild-type female mosquitoes when contamination was absent. In contrast, the perfect eradication policy failed easily even if a small percentage of *Wolbachia*-infected female mosquitoes are contaminated by the release of *Wolbachia*-infected male mosquitoes, whereas a decrease in the total number of female mosquitoes at a certain level is possible. Our study also found that *Wolbachia*-infected male mosquitoes need not be released until the female mosquitoes become extinct and there exists an optimal stopping point of the release at which we can obtain sufficiently successful control. This result suggests a new insight into reducing the economic burden when it comes to actual policy implementation. However, our study also suggests that we need a different policy under the scenario of contamination. In this scenario, the release stop strategy has little effect on reducing the release costs. Furthermore, the release cost was determined by the scale of the release rate.

Similarly with the previous field work (Zheng et al. 2019), our modeling study also suggested that IIT cannot be an effective method to eradicate female mosquitoes perfectly when contamination is present. However, our study using the Malaria-IIT model found that female mosquitoes need not be eradicated perfectly to extinguish malaria and we can obtain a successful control result even with contamination occurring. We also found that there exists an optimal release rate at which the release cost becomes minimal, which should be smaller when the contamination ratio is higher. On one hand, the extinction time is prolonged when we choose a lower release rate. Therefore, a measure should be carefully chosen which is optimal in designing a control strategy for preventing mosquito-borne diseases.

Every year, many people are infected and lose their lives because of vector-borne diseases. However, it is difficult to implement a control policy because field experiments are spatially and temporally restricted. In particular, the IIT method has been shunned owing to contamination. Our study suggests that the IIT method is very promising, even though perfect eradication is not possible in real and it might be valuable to be tested in the real field of endemic. In this study, we did not consider spatial effects, such as geographical conditions and spatial spreading, which we consider as a future work.

## Data availability

Data and codes are available on request.

## Acknowledgements

This work was partially supported by a Grant-in-Aid for Scientific Research (KAKENHI, JP17KK0094), and Japan Science and Technology Agency(JST) CREST to S.S.L. (JPMJCR2111).

## Supplementary

### A Equilibria and liner stability of wild-type mosquitoes

The equilibria (*F*\*, *M*\*) of the wild-type mosquitoes satisfy the equations;

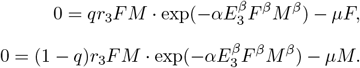

With taking *q* = 0.5, we have *F*\* = *M*\* and

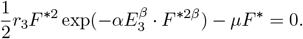

Next, we analyse the stability of the equilibrium. With setting

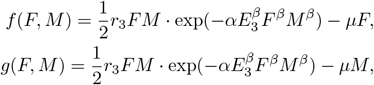

we obtain the Jacobian matrix

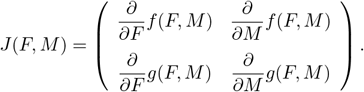

and the eigenvalue equation of *λ*^2^ − (*λ*_1_ + *λ*_2_)*λ* + *λ*_1_ · *λ*_2_ = 0, where

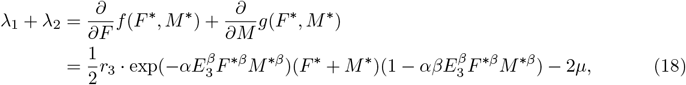

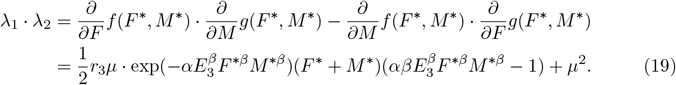

We numerically evaluated the stability by judging the sign of the equations (18) and (19) with given parameter values.

